# Extracellular signalling modulates Scar/WAVE complex activity through Abi phosphorylation

**DOI:** 10.1101/2021.10.09.462405

**Authors:** Shashi Prakash Singh, Peter A. Thomason, Robert H. Insall

## Abstract

The lamellipodia and pseudopodia of migrating cells are produced and maintained by the Scar/WAVE complex. Thus, actin-based cell migration is largely controlled through regulation of Scar/WAVE. Here we report that the Abi subunit - not Scar/WAVE – is phosphorylated in response to extracellular signalling. Like Scar, Abi is phosphorylated after the complex has been activated, implying that Abi phosphorylation modulates pseudopodia, rather than causing new ones to be made. Consistent with this, Scar/WAVE complex mutants that cannot bind Rac are also not phosphorylated. Several environmental cues also affect Abi phosphorylation - cell-substrate adhesion promotes it and increased extracellular osmolarity diminishes it. Both unphosphorylatable and phosphomimetic Abi efficiently rescue the chemotaxis of Abi KO cells and pseudopodia formation, confirming that Abi phosphorylation is not required for activation or inactivation of the Scar/WAVE complex. However, pseudopodia and Scar/WAVE patches in the cells with unphosphorylatable Abi protrude for longer, altering pseudopod dynamics and cell speed. Cells in which Scar and Abi are both unphosphorylatable can still form pseudopods, but migrate substantially faster. We conclude that extracellular signals and environmental responses modulate cell migration by tuning the behaviour of the Scar/WAVE complex after it has been activated.

## 1. Introduction

In migrating cells, the Scar/WAVE complex acts as a principal catalyst to generate actin rich protrusions. In particular, lamellipods or pseudopods are generated by activation of the Arp2/3 complex controlled by the Scar/WAVE complex, leading in turn to polymerization and growth of actin filaments [1]. The five-membered complex consists of Pir121, Nap1, Scar/WAVE, Abi and HSPC300 subunits [2]. It is regulated by the small GTPase Rac1. Rac1 does not interact with the complex when GDP-bound, but when GTP-bound it interacts with the Pir121 subunit, making the Scar/WAVE complex active and able to promote actin polymerization [3, 4]. Multiple other regulators affect the Scar/WAVE complex’s activity [1].

Several authors have reported that phosphorylation, of either the Scar/WAVE [5–10] and Abi [7, 11, 12] subunits, is a necessary and important activator of the complex and for the formation of proper lamellipodia or pseudopodia. However, we recently showed that Scar/WAVE phosphorylation is not required for the Scar/WAVE complex to be activated in either mammalian or *Dictyostelium* cells; rather, it modulates pseudopod dynamics [13]. In this work we investigate if the same is true for Abi.

Several authors have reported phosphorylation of Abi [6, 7, 11, 12, 14, 16]. Phosphorylation by the tyrosine kinase Abl (Abelson kinase) has been shown to control the stability of Abi, as well actin morphology, in both blood cells [11, 14] and in *Drosophila melanogaster* [12]. Other reports describe ERK2 phosphorylation sites in Abi [7, 16], and suggest that they activate protrusion and control its dynamics during migration, though we note that our recent data strongly oppose the previously described role for ERK2 in Scar/WAVE activation. CDK1-mediated serine phosphorylation in Abi may be important in mitosis [17].

ERK2 and Abl have specific recognition sequences, which are not conserved in Abi proteins from different eukaryotic organisms. Moreover, receptor tyrosine kinases are not present in *Dictyostelium* [15], despite strong conservation in the sequence and activation mechanisms of its Scar/WAVE complex. We have therefore investigated the importance of Abi phosphorylation in *Dictyostelium* cell motility.

## 2. Materials and Methods

### 2.1 Dictyostelium cell culture

Wild type strains of *Dictyostelium discoideum*, Ax3 and NC4, were obtained from the *Dictyostelium* Stock Center (http://dictybase.org/StockCenter) [18]. All cell lines used and generated in this study are mentioned in Table S1. Axenic culture of cells was maintained in HL5 medium (Formedium) supplemented with 100 units of penicillin and 100 mg/ml streptomycin-sulfate in Petri dishes or in shaken suspension at 2-4×10^6^ cells/ml (22°C, 125 rpm). Non-axenic culture of cells was maintained on SM agar (Formedium) plates with *Klebsiella aerogenes*. Cells were mixed in *Klebsiella aerogenes*, spread on SM agar plates with a sterile L-rod spreader and kept in a 22°C incubator for approximately 40 h to form a clearing zone enriched with amoebal cells.

### 2.2 Plasmid constructs

A shuttle vector containing the *abi* gene was constructed by excising *abi* from vector pAD48 [19] with BglII-SpeI and cloning it into pDM344. To replace the actin15 promoter from the resulting vector, the 5’UTR of *abi* was PCR-amplified from *Dictyostelium discoideum* genomic DNA and cloned using EcoRI-ClaI. The NgoMIV fragment from the above construct was subcloned into pDM304 and pDM459 to generate Abi and HSPC300-eGFP co-expression constructs. Phosphorylation mutants of Abi were generated by PCR-based site-directed mutagenesis using primers listed in Table S2. To construct Pir121-EGFP knock-in constructs, the 3’ UTR of Pir121 was PCR amplified from genomic DNA using the primers listed in Table S2 and cloned into pPT808 [4] using Gibson assembly (NEB). All plasmids used in this study are listed in Table S3.

### 2.3 Generation of Scar-/Abi-double knock out cells

The *scar*-/*abi*- double knockout was generated by knocking out *abi* in *scar*- cells (strain IR46) by homologous recombination using knock out vector pAY16P[20]. Briefly Scar- cells were transfected with 10 μg KpnI-MluI linearized pAY16P. 24 h after transfection, *scar-/abi-* knockout clones were selected with 10 μg/ml blasticidin in 96 well plates. Knockout clones were confirmed by PCR using *abi* UTR and blasticidin primers mentioned in Table S2, and western blotting.

### 2.4 Generation of Scar-/Abi-/pir121-eGFP cells

The Pir121-eGFP knock in construct was generated by tranfecting *scar-/abi-* cells with a gel purified *pir121eGFP* region, excised with EcoRI. eGFP -labelled cells were FACS-sorted 48h post transfection. Positive knock in clones were confirmed by western blotting and AiryScan confocal microscopy.

### 2.5 Transfection of Dictyostelium cells

To transfect using extrachromosomal plasmids, 1 ×10^7^ cells/ml were washed once with electroporation buffer and resuspended in 420 μl EB (EB; 5mM Na_2_HPO_4_, 5mM KH_2_PO_4_ and 50mM sucrose) and approximately 0.5 μg plasmid DNA was mixed with cells in 2mm gap electroporation cuvettes, and electroporated into the cells by pulsing once at 500V using an ECM399 electroporator (BTX Harvard). Cells were then transferred into HL5 medium. After 24 h, 10 μg/ml G418, blasticidin or 50 μg/ml hygromycin were added to select transformants. To make knockout or knockin clones, 10 μg linearized vectors were electroporated into cells.

### 2.6 GFP-TRAP pulldown

Cells grown in a 15 cm dish were lysed with 1ml TNE/T buffer (10 mM Tris-HCl pH 7.5, 150 mM NaCl, 0.5 mM EDTA and 0.1 % Triton X-100) containing HALT protease and phosphatase inhibitors. Lysates were kept on ice for 5 mins and cleared by centrifugation (13000 rpm, 4 °C, 5 mins). 25 μL GFP-TRAP beads (ChromoTek) were washed twice with TNE/T buffer and resuspended in 100 μL TNE/T buffer. 1 μg of lysate was added to the beads and kept on rotation for 30 min at 4 °C. Beads were spun down at (2700g, 4 °C, 2 mins) and washed 3 times with TNE/T buffer. Proteins from the beads were eluted by adding 50 μL 2 x NuPAGE LDS sample buffer and boiled (100 °C, 5 mins).

### 2.7 Western blotting

Cells were lysed by directly adding NuPAGE LDS sample buffer (Invitrogen) containing 20mM DTT, HALT protease and phosphatase inhibitors (Thermo Fisher Scientific) on top of cells and boiled at 100 °C for 5 mins. Proteins were separated on 10 % Bis-Tris NuPAGE gels (Invitrogen) or on hand-poured low-bis acrylamide (0.06% bis acrylamide and 10% acrylamide) gels, then separated at 150 V for 90 mins. Proteins were transferred onto 0.45μM nitrocellulose membrane. Membranes were blocked in TBS+5% non-fat milk. Primary antibodies were used at 1:1000 dilution. 1:10000 fluorescently conjugated secondary antibody was used to detect the protein bands by Odyssey CLx Imaging system (LI-COR Biosciences). Mccc1 was used as loading control [21].

### 2.8 Phosphatase treatment

Cells grown in a 35 mm Petri dish were lysed in 100 μL TN/T buffer (10 mM Tris-HCl pH 7.5, 150 mM NaCl and 0.1 % Triton X-100), kept on ice for 5 mins and cleared by centrifugation (13000 rpm, 4 °C, 5 mins). Proteins were dephosphorylated using 1μL Lambda phosphatase at 30 °C (NEB; P0753S) for 1 h. Protein dephosphorylation was assessed by western blotting using low-bis gels.

### 2.9 Under agarose chemotaxis

Cellular morphology, pseudopod dynamics and cell migration were measured by an under-agarose folate chemotaxis assay as described earlier [13]. In brief, 0.4% SeamKem GTG agarose was dissolved in boiling Lo-Flo medium (Formedium). After cooling, 10 μM folic acid was added. 5 ml of agarose-folate mix was poured into the 1% BSA-coated 50mm glass bottom dishes (MatTek). A 5 mm wide trough was cut with sterile scalpel and filled with 200 μl of 2× 10^6^ cells/ml. Cell migration was imaged after 4-6h with 10x and 60x DIC. To examine the localization of labelled proteins in the pseudopods, cells were also imaged using an AiryScan confocal microscope (Zeiss).

### 2.10 Stimulation of cells with folate and cAMP

NC4 or Ax3 cells were grown non-axenically and washed 3 times with KK2 buffer, resuspended at 2×10^7^ cells/ml and incubated for 1h (125 rpm, 22 °C). Cells were stimulated with 200μM folate and lysed in LDS sample buffer at 0, 5, 10, 20, 30 45, 60, 90, 120 secs and tested by western blotting.

For cAMP stimulation, 2×10^7^ cells/ml cells in KK2 were starved. After 1h, cells were pulsed with 100 nM cAMP every 6 min for 4 h. 2mM caffeine was added to inhibit internal cAMP signalling. After 1 min of caffeine addition cells were stimulated with 10μM of cAMP and lysed in 1X LDS sample buffer at 0, 5, 10, 20, 30 45, 60, 90, 120 secs and used for western blotting.

### 2.11 Microscopy

To determine the morphology, chemotactic speed and directionality of cells phase-contrast time lapse microscopy was performed at 10×/0.3NA on a Nikon ECLIPSE TE-2000-R inverted microscope equipped with a Retiga EXI CCD monochromatic camera. Images of cells migrating under agarose up a folate gradient were captured every? minute for 45 minutes. DIC images were taken every 2 s for 3 minutes with 60x/1.4 NA to observe pseudopod formation. HSPC300-eGFP expressing cells were used to determine the activation of the Scar complex. The localization of Scar/WAVE complex, Arp2/3 complex, and F-Actin was examined by a 63x/1.4 NA objective on an AiryScan Zeiss 880 inverted confocal microscope.

### 2.12 Quantification of data and statistics

Every experiment was performed at least three times. Speeds of cells were calculated using a homemade plugin in ImageJ. Pseudopodia and Scar/WAVE complex dynamics were calculated by counting the number of frames during a pseudopodia extension manually using ImageJ. Graphs and statistical analyses were performed using Prism.

To quantify the proportion of Abi and Scar/WAVE phosphorylation, the total intensity of all bands and the lowest band were determined using the Odyssey CLx Imaging system (LI-COR Biosciences). The percentage of phosphorylation was calculated by using the formula: % phosphorylation=(Total intensity-intensity of lowest band)*100/Total intensity.

Statistical significance analyses were performed by non-parametric statistics, such as one-way ANOVA and Dunn’s multiple comparison tests, as described for each result. The experimental groups were tested for normal distribution using the Shapiro-walk test of Prism 7 software (Graphpad, La Jolla, USA). Sample sizes are provided in the figure legends. n refers to; independently repeated experiments in western blotting, total number of cells from 3 independent experiments for cell speed, pseudopod dynamics, and Scar patch frequency, duration of patches/accumulation size in ≥25 or more cells.

## 3. Results

### 3.1 Constitutive phosphorylation of Abi

Analysis of Abi phosphorylation is difficult with standard techniques – normal PAGE lacks resolution, while mass spectrometry detects phosphosites in polyproline domains poorly. In previous studies of Scar/WAVE, we have successfully used low-bis acrylamide SDS-PAGE and western blotting [13, 22]. In normal PAGE gels, Abi from migrating *Dictyostelium* cells runs as a single band (Figure 1A), with quantitative analysis showing this band forming a single diffuse peak (Figure 1B). However, low-bis acrylamide gels resolve identical samples of Abi into two distinct bands (Figure 1C) with clear separation apparent in intensity plots (Figure 1D). The two bands resolve into one upon phosphatase treatment (Figure 1E), and two intensity peaks (solid line; Figure 1F) shift to a single peak (dotted line; Figure 1F), confirming both that the multiple bands are due to phosphorylation, and that Abi is phosphorylated in unstimulated cells.

**Figure 1.**
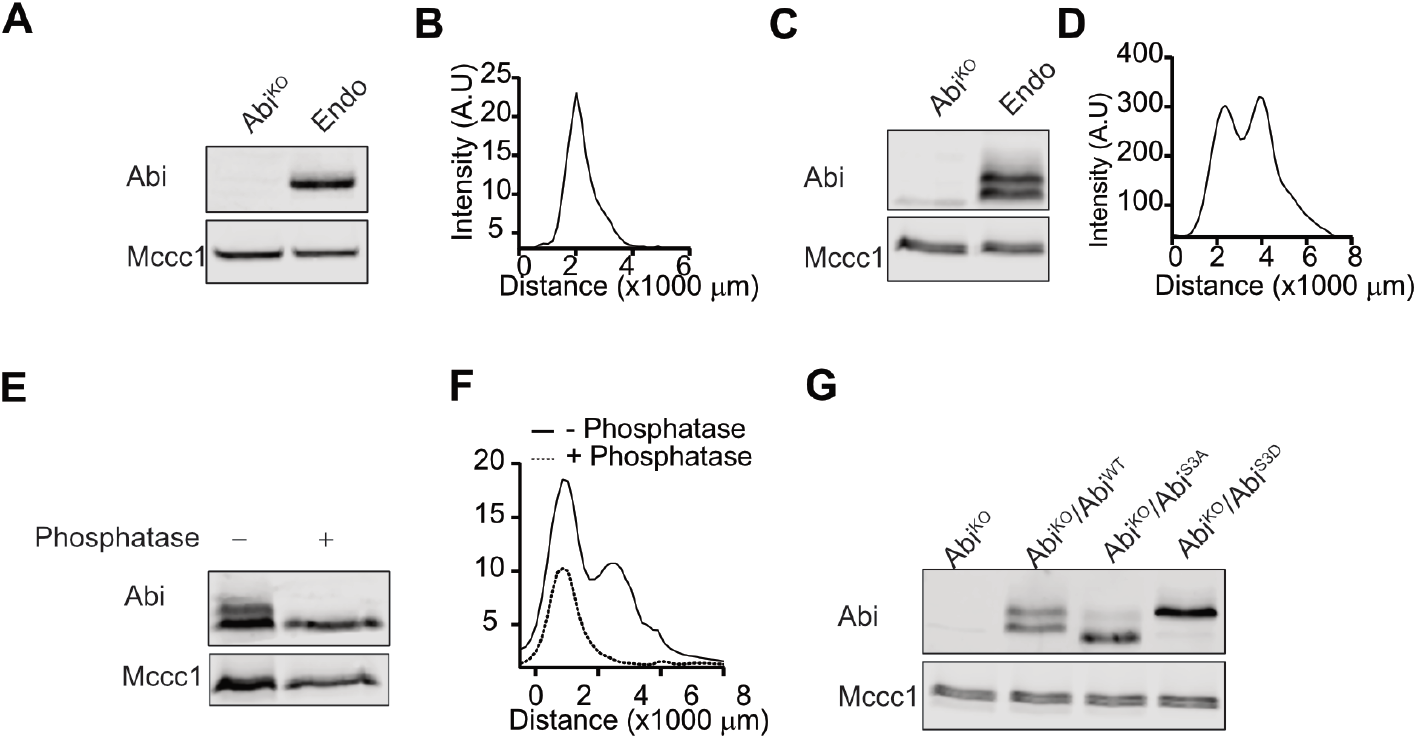
Multiple phosphorylations in Abi. (A) Western blot of *Dictyostelium* Abi using normal (0.3%) bis-acrylamide gels. Whole cells were lysed in sample buffer, boiled, and separated on 10 % bis-tris SDS-PAGE gels, blotted, and probed with an anti-Abi antibody, yielding a single band. Mccc1 is a loading control. B) Line scan of Abi band signal intensity shows a single peak. (C) Western blot of Abi using low (0.06%) bis-acrylamide gels, yielding two discrete bands. (D) Line scan of Abi bands’ signal intensities shows separate peaks with broad base. (E, F) Phosphatase treatment. Whole *Dictyostelium* cells were lysed in TN/T buffer, then incubated with and without lambda phosphatase, boiled in sample buffer, and analyzed using low- Bis acrylamide gels. The scan shows two peaks resolved into one and shifted downwards after phosphatase treatment. (G) Unphosphorylatable and phosphomimetic triple mutations of serines S166/168/169 analysed by western blotting on low-bis gels. Unphosphorylatable mutants run as a single, high-mobility band; phosphomimetic ones as a single low-mobility band. KO; knock-out, Endo; endogenous.

Previous identification of Abi phosphorylation sites relies on prediction, *in vitro* phosphorylation assays [7, 16] or overexpression of tagged proteins [12]. Overexpression of the Scar/WAVE subunits can result in subunits that are not incorporated into a full complex, with strong potential for artifacts. To avoid these secondary effects of overexpression, we explored phosphorylation in native Abi. Mass spectrometry analysis of the full complex, immunoprecipitated using GFP-TRAP on cells with stably tagged GFP-Nap1, did not identify any phosphosites in Abi. Given the clear phosphorylation seen in Fig. 1 this could be due to two reasons - either the phosphosites were inaccessible to trypsin or chymotrypsin cleavage, or they could not be detected by mass spectrometry (a particular problem for phosphosites in the polyproline domain). We therefore screened the plausible serines (S) and threonines (T) in Abi by mutating them to unphosphorylatable alanines (S/T➔A) and phosphomimetic aspartates (S/T➔D). This led us to focus on three serines - S166,168,169. Simultaneous mutation of all three residues to unphosphorylatable (Abi^S3A^; S166/168/169A) forms and expression in Abi^KO^ cells showed near-total loss of the additional band of Abi (Figure 1G). A trace band is just visible, implying a minimal level of phosphorylation at another site. Similarly, phosphomimetic mutation of all three serines to phosphoforms (Abi^S3D^; S166/168/169D) caused constitutive mobility shift on the gel (Figure 1G). This further confirms that the additional band of Abi is due to phosphorylation.

### 3.2 Chemoattractant stimulation of Abi Phosphorylation

Previous studies have shown that extracellular signalling causes cytoskeletal changes and increases protrusions [23–25]. This is presumably mediated in large part through the Scar/WAVE complex. We have previously shown that phosphorylation of the Scar/WAVE subunit does not activate the complex, nor is it induced by chemoattractant signaling [13], and that the constitutive phosphorylation on the C-terminus of Scar/WAVE makes it less active [26]. In mammalian cells, Abi phosphorylation is thought to be mediated by nonreceptor tyrosine kinase Abl and MAP kinase signalling [7, 12, 14]. *Dictyostelium* lacks nonreceptor tyrosine kinases of the Abl & Src family, and none of the phosphosites of Abi fit the MAP kinase consensus sequence (S/TP). Hence, we examined the changes in the Scar/WAVE redistribution and Scar and Abi phosphorylation in response to chemoattractant stimulus. *Dictyostelium* uses two principal chemoattractants - folate and cAMP, during growth and development respectively. Treatment of growing cells with folate and developed cells with cAMP causes a rapid change in the amount of polymerized actin [27, 28]. Brief exposure of cAMP can also cause rapid recruitment of the Scar/WAVE complex to the cell periphery [29].

To determine the correlation between changes in the Scar/WAVE complex recruitment and phosphorylation in response to chemoattractant stimulus, we examined Scar/WAVE recruitment by AiryScan confocal microscope imaging, and phosphorylation of Scar and Abi by western blotting. Active Scar/WAVE complex accumulates in the pseudopods as a short-lived bright patch or (occasionally) flash. Soon after folate treatment of *Dictyostelium* cells expressing GFP-Nap1[26], the Scar/WAVE redistributes to the cell periphery as puncta, and remains in this state for at least 60 seconds (Figure 2A, Video S1). This correlated with a substantial increase in Abi phosphorylation after folate treatment, without significant changes to Scar (Figure 2B, C&D). In the same way, cAMP stimulation of chemotactically competent cells causes similar redistribution of Scar/WAVE (Figure 2E) and increased phosphorylation of Abi (Figure 2F, G&H, Video S2). A small minority of Scar was also phosphorylated in response to cAMP. These results raised the possibility that Abi, but not Scar, phosphorylation couples chemoattractant signalling to the actin cytoskeleton.

**Figure 2.**
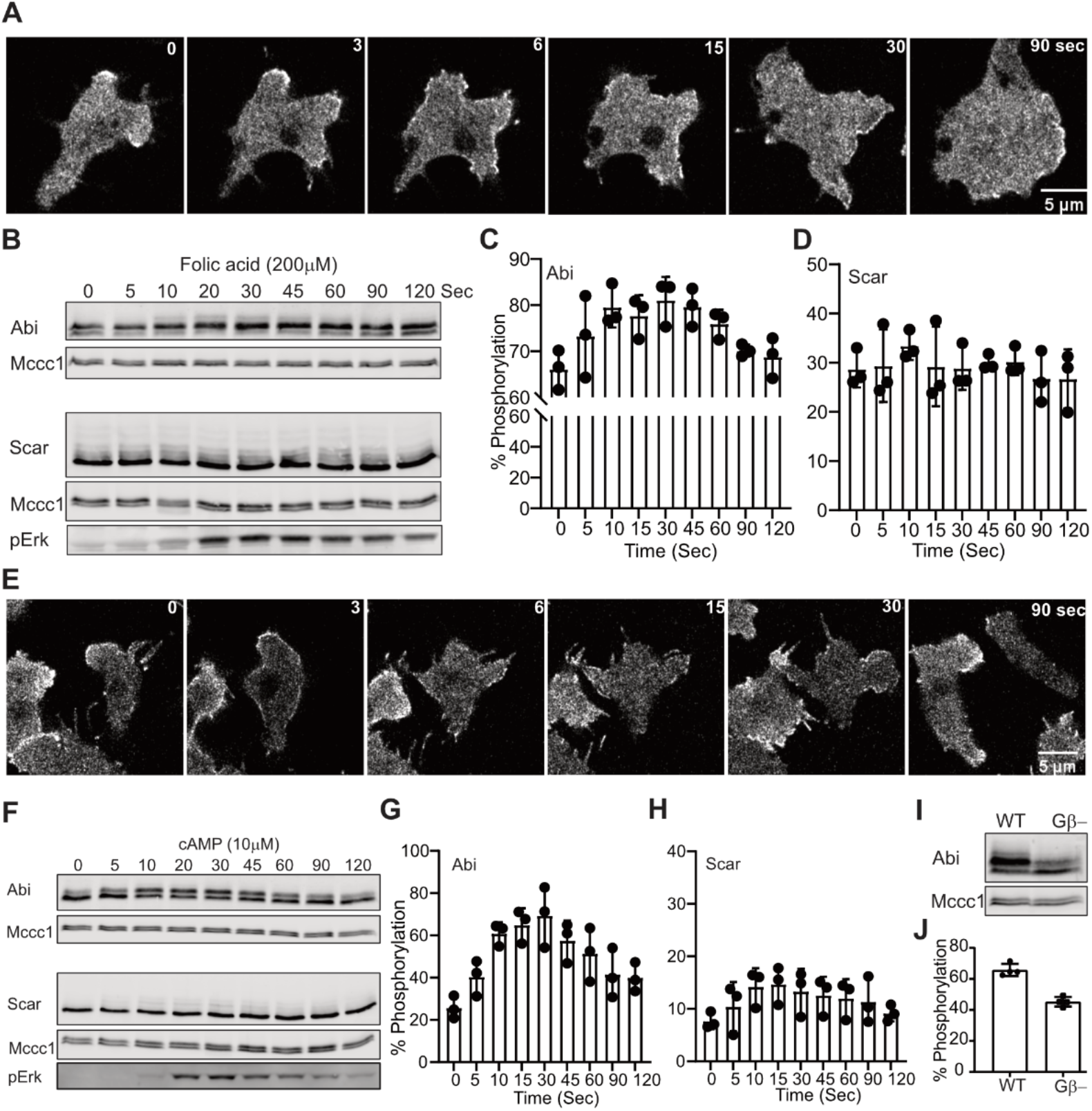
Signaling dependent phosphorylation of Abi and Scar. (A) Redistribution of the Scar/WAVE complex after folate treatment. *Dictyostelium* cells expressing EGFP-Nap1, which labels the Scar/WAVE complex, were imaged by AiryScan confocal microscopy. The Scar/WAVE complex rapidly gets redistributed as puncta on the cell membrane periphery after addition of folate (200 mM) treatment (Scale=5 mm). (B-D) Effect of folate stimulation on Abi and Scar phosphorylation. Growing *Dictyostelium* cells were washed and treated with 200 mM folate for the indicated time points, then Abi and Scar band shifts were analyzed using low Bis-acrylamyde gels. Folate enhances Abi phosphorylation (C; mean±SD, n=3) but not Scar phosphorylation (D; mean±SD, n=3) despite stimulation of signaling shown by Erk phosphorylation. (E) Redistribution of the Scar/WAVE complex after cAMP treatment. cAMP competent *Dictyostelium* cells expressing EGFP-Nap1 were imaged by AiryScan confocal microscopy. The Scar/WAVE complex gets redistributed rapidly on the cell membrane periphery after addition of cAMP (10 mM) treatment (Scale 5 mm). (F-H) Effect of cAMP stimulation on Abi and Scar phosphorylation. cAMP competent cells were stimulated with 10 mM cAMP for the indicated time points, then Abi and Scar band shifts were analyzed by western blotting. cAMP enhances Abi phosphorylation (G; mean±SD, n=3) substantially with little change in Scar phosphorylation (H; mean±SD, n=3) despite huge activation of signalling shown by Erk phosphorylation. (I-J) Abi phosphorylation in WT and Gβ- cells. Abi band shifts were analyzed in WT and Gβ- cell by western blotting. Abi in Gβ- cells has substantially less phosphorylation (J; mean ± SD; n = 4). cAMP, cyclic 3’,5’-adenosine monophosphate; ERK, extracellular signal regulated kinase; Gβ, G-protein beta subunit; WT, wild type.

To further support this finding we measured Abi phosphorylation in signalling deficient G-protein beta knockout (Gβ-) cells [30]. As expected, Abi phosphorylation was greatly reduced in Gβ- cells (Figure 2I&J). Thus, chemotactic signalling stimulates an increase in Abi phosphorylation.

### 3.3 Cell-substrate adhesion and osmotic shock alter Abi phosphorylation

Cell-substrate adhesion is an important regulator of pseudopods and lamellipod dynamics. We recently showed that cell-substrate adhesion increases Scar phosphorylation [13, 22]. We therefore examined the role of cell-substrate adhesion on Abi phosphorylation, exploiting the ability of *Dictyostelium* to grow in suspension and adhesion. First, we allowed suspended cells to adhere to a Petri dish and followed changes in Abi phosphorylation by western blotting. This revealed an obvious (and significant - p=0.0082 Kruskal–Wallis test) increase in Abi phosphorylation in adhered, compared to suspended, cells (Figure 3A&B). The phosphorylated band fraction increased from 39.7±12.3% to 66±8.9% (mean±SD; Figure 3B). In contrast, when we de-adhered cells using a stream of growth medium from a pipette, we observed a clear drop in the relative abundance of the phosphorylated upper band (Figure 3C). The phosphorylated Abi dropped from 50.3±3.9% to 33.8±2.4% of the total (mean±SD; Figure 3D). Thus, as with Scar phosphorylation, cell-substrate adhesion induces Abi phosphorylation.

**Figure 3.**
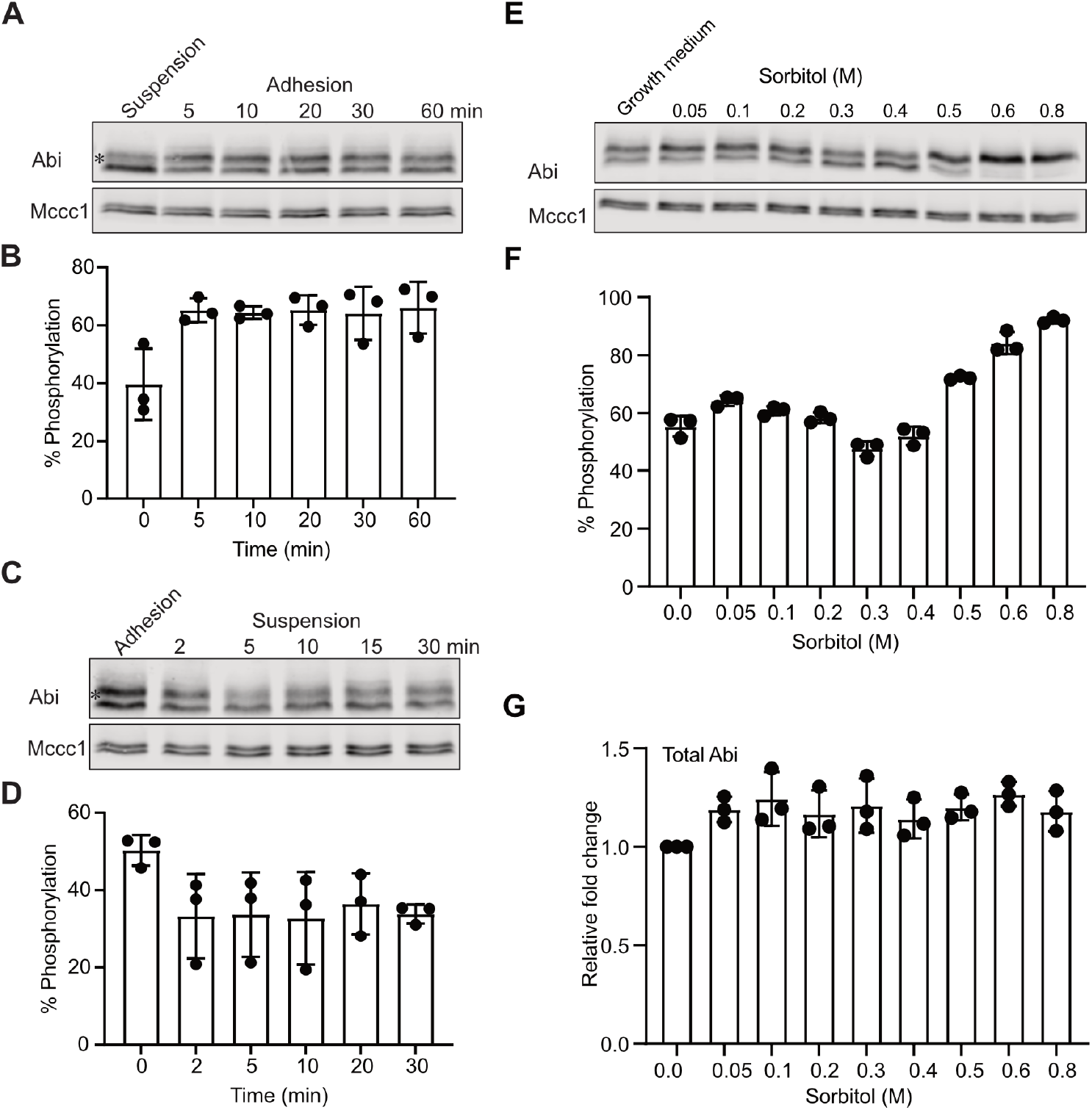
Cell-substrate adhesion enhances Abi phosphorylation. (A-B) Effect of adhesion on Abi phosphorylation. Suspension grown cells were allowed to adhere in Petri dishes, lysed at indicated time points, and cell lysates were analyzed by western blotting. The intensity of upper Abi band (*) increases after adhesion. Quantitation of lane band intensities (intensity of “upper bands”/total band intensity; mean ± SD, n = 3) confirms increased intensities of phosphorylated Abi bands after adhesion (B). (C-D) Effect of de-adhesion on Abi phosphorylation. Cells grown in adhesion on Petri dishes were detached by a jet of medium from a 10 ml pipette, collected and maintained in suspension (approx. 2×10^6^ cells/ml) by shaking at 120 rpm. Cell lysates were analyzed by western blotting. Quantitation of lane intensities (mean ± SD, n = 3) confirms a loss of phosphorylated Abi bands (*) intensity(D). (E-G) Effect of various sorbitol concentrations on Abi and Scar phosphorylation. Medium from cells was replaced with PB containing various concentrations of sorbitol. After 5 mins cell lysates were prepared in sample buffer and analyzed for Abi (D) band shifts. Abi phosphorylation is abolished at 400 mM sorbitol and induced with >400 mM sorbitol (F; mean ± SD, n = 3) without affecting total Abi concentration (G; mean ± SD, n = 3).

*Dictyostelium*, like other chemotactic cells, change shape and reorganize their actin cytoskeleton upon stimulus by chemoattractant. Likewise, cells also cope with the hyper-osmotic stress by re-organizing F-actin and changing their shape [31–33]. Additional proteins that translocate to the cell cortex in response to the hyperosmotic stress are myosin [34], S-adenosyl-L-homocysteine hydrolase (SAHH) and cofilin [31]. A correlation between hyper-osmotic stress, crosslinking of α-actinin and gelation factor with F-actin has been observed earlier [32]. In cell protrusions, F-actin polymerization is principally catalyzed by the Scar/WAVE complex, but the effect of osmotic stress is unknown. We examined live cell shape changes by DIC microscopy, and the recruitment of the Scar/WAVE complex and F-actin distribution by AiryScan confocal imaging after adding sorbitol (0.4M) to cells attached to a glass surface. Cells retracted their protrusions, stopped movement, and shrunk quickly after addition of sorbitol (Figure S1A, Video S3). The Scar/WAVE complex from the pseudopods (arrow; Figure S1B, Panel1), translocated to the cell cortex upon sorbitol treatment (Figure S1B, Video S4). F-actin localization was initially mainly in the pseudopod (arrow; Figure S1C, panel 1), but redistributed not only to the cell cortex, but also substantially to large vesicles, presumably internalised macropinosomes (asterisks; Figure S1C, Video S5).

Redistribution of the Scar/WAVE complex from the pseudopods to cell cortex suggests changes in its activity (Figure S1B) in response to hyper-osmotic stress. We therefore examined Abi phosphorylation after cells were treated with various concentrations of sorbitol. Phosphorylation of Abi was unaffected by low concentrations of sorbitol (0.1M), but was somewhat inhibited by high concentrations (≤0.4M) (Figure 3E&F). High concentrations of sorbitol (0.5-0.8M) induced Abi phosphorylation (Figure 3E&F) without affecting total Abi (Figure 3G). This result shows that Abi phosphorylation is affected by multiple changes in the extracellular environment.

### 3.4 Abi phosphorylation is activation-dependent

The phosphorylation of Scar and Abi are often reported as events upstream of Scar/WAVE complex activation [7]. That is, most authors assert that Scar/WAVE complex members are phosphorylated in order to make them active, or to increase the probability that they will be activated. However, we recently found that phosphorylation of Scar is a consequence of the complex activation and not a cause [13]. This has important implications for the mechanism of activation - it implies that phosphorylation’s role is unimportant to the mechanism of complex activation. To investigate whether Abi phosphorylation is a post-activation event, like Scar/WAVE, we tested the effect of latrunculin A (an inhibitor of actin polymerization [35, 36]) on Scar/WAVE recruitment and Abi phosphorylation. Cells treated with latrunculin, but no chemoattractant or upstream signal, showed exaggerated recruitment of the Scar/WAVE complex and hyperactivation of the Arp2/3 complex (Figure 4A, Video S6) along with induction of Scar phosphorylation in *Dictyostelium* cells [13, 37]. As with Scar [22], Abi phosphorylation increased rapidly upon latrunculin treatment (Figure 4B), with similar kinetics to the increases in Scar and Arp2/3 recruitment. The fraction of Abi that was phosphorylated increased (Figure 4C), with a slight reduction in total Abi upon latrunculin A treatment (Figure 4D). This suggests that Abi phosphorylation is driven by Scar/WAVE activation, rather than upstream signalling.

**Figure 4.**
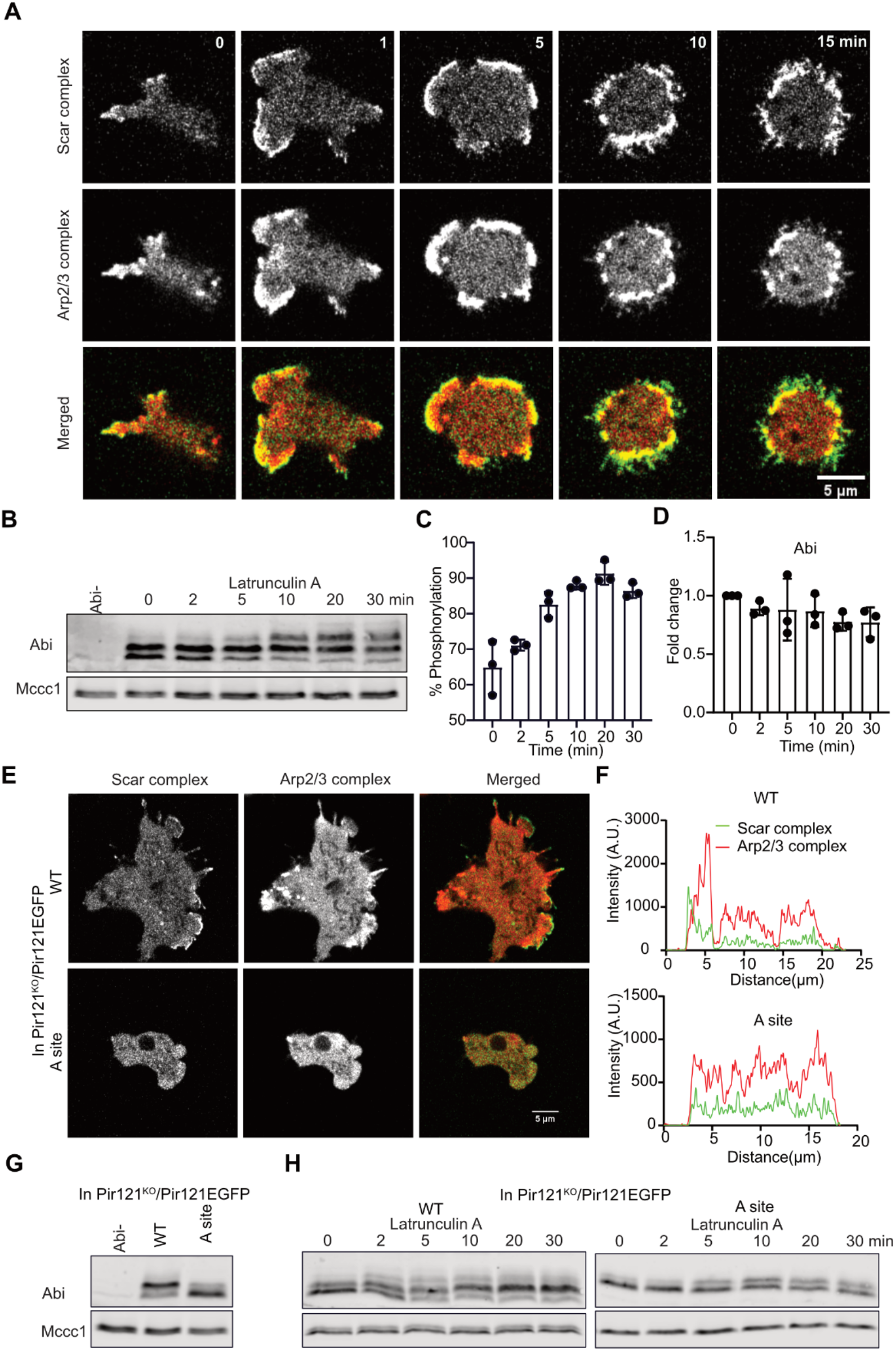
Effect of the activation of the Scar/WAVE complex on Abi phosphorylation. (A) Hyperactivation and redistribution of the Scar/WAVE complex after latrunculin A treatment. Cells expressing EGFP-Nap1 and mRFP-mars2-ArpC4 were imaged using an AiryScan confocal microscope. Latrunculin A (5 μM) was added at t = 30 seconds. Scale bar = 5 μm. Addition of latrunculin enhances the Scar complex and Arp2/3 recruitment at the cell periphery. (B-D) Increased Abi phosphorylation after latrunculin A treatment. Cells treated with latrunculin A for the indicated times were lysed in sample buffer, then Abi band shift and intensities were analyzed by western blotting. Phosphorylated Abi bands become more abundant due to latrunculin A treatment (C; mean±SD, n=3), with a slight reduction in total Abi (D; mean±SD, n=3). (E) Importance of A-site in the activation of the Scar/WAVE complex and Arp2/3 complex. WT and the A-site mutant of Pir121 were co-expressed with mRFP-mars2-ArpC4 in the Pir121- cells and imaged by AiryScan confocal microscopy under agarose chemotaxis up a folate gradient. The A-site mutant is unable to activate the Scar/WAVE and Arp2/3 complexes. (F) Quantitation of Scar and Arp2/3 activation in WT and A-site mutant from merged panels. A line was drawn across the centre of the cell and relative pixel values plotted. (G) Importance of Rac1-binding PIR121 A site for Abi phosphorylation. WT, A site (K193D/R194D) Pir121 expressing cells were analyzed for Abi band shifts by western blotting. Phosphorylated Abi bands are absent in A site mutant. (H) Effect of latrunculin A treatment on Abi phosphorylation in WT and A-site mutant. WT and A site mutant of Pir121 were treated with latrunculin A for the indicated times, lysed in sample buffer, then Abi band shift and intensities were analyzed by western blotting. Phosphorylated Abi bands become abundant in WT cells after latrunculin A treatment. A.U. arbitrary unit, WT, wild type, A-site; adjacent site.

Scar/WAVE complex activation completely depends on GTP-bound Rac1 interacting with the Pir121 subunit [3, 38, 39]. We therefore examined Abi phosphorylation in cells with the A-site mutation in Pir121, which cannot bind Rac1 [3, 4]. This makes the cells excellent for testing whether phosphorylation is upstream or downstream of activation - signalling pathways upstream will be unaffected, whereas activationdependent processes will be lost. As expected, A-site mutants activated neither the Scar/WAVE nor the Arp2/3 complex (Figure 4 E&F), confirming the inactive state of the Scar/WAVE complex. Abi phosphorylation was abolished in the A-site mutant (Figure 4G). Furthermore, treatment with latrunculin A did not stimulate Abi phosphorylation in A-site mutant cells (Figure 4H). Altogether, these results confirm that Abi phosphorylation is a result of Scar/WAVE complex activation.

### 3.5 Abi phosphorylation tunes cell migration and pseudopod formation

In normal cells, every lamellipod or pseudopod’s Arp2/3-mediated actin polymerization is driven by the Scar/WAVE complex[2]. There have been several analyses of phosphorylation’s roles. Tyrosine phosphorylation in Abi has been implicated in the proper localization of the Scar/WAVE complex formation in *Drosophila* [12]. Multiple growth factor-stimulated and Erk-dependent phosphorylation sites (S183, 216, 225, 392, 410 and T265, 267, 394) have been reported to increase Scar/WAVE’s interaction with the Arp2/3 complex [6, 16]. However, all these studies were done with either fluorescently-tagged, overexpressed or bacterially-expressed Abi *in vitro*, meaning they are at risk of being nonphysiological. To observe the effects of Abi phosphorylation as part of the normal Scar/WAVE complex in moving cells, we co-expressed unlabelled Abi and phospho-mutants (AbiS3A and AbiS3D) and HSCP-300-eGFP [40] in Abi null (Abi^KO^) cells. Both of the phospho-mutants were expressed at the normal level. Western blotting of Pir121, Nap1, Scar and Abi from pull down samples showed that Abi^WT^, Abi^S3A^ and Abi^S3D^ formed stable complexes with them (Figure S2A, B&C). To analyze the migratory phenotype, we examined cells migrating under agarose up a folate gradient. Abi^WT^, Abi^S3A^ and Abi^S3D^ all localized at the pseudopodia of cells, with marked differences in dynamics (Figure 5A, Video S7). The localized bursts (Scar patches) of Abi^WT^ recruitment lasted 11.8±4.6 seconds (mean±SD), and the phosphorylation-deficient Abi (Abi^S3A^) remained at the pseudopod edges for 12.3±5.3 seconds (Mean±SD) (Figure 5B). However, the Abi^S3D^ recruitment was short-lived (8.2±2.6 seconds, mean±SD)(Figure 5B) and Abi^S3D^ patches were smallest at the accumulation site (Figure 5C). This led to changes in the patch frequency- the smaller and shortest-lived Abi^S3D^ patches were made at a higher frequency than Abi^WT^ and Abi^S3A^ (Figure 5D). These results imply that Abi phosphorylation modulates the dynamics of the Scar/WAVE patches.

**Figure 5.**
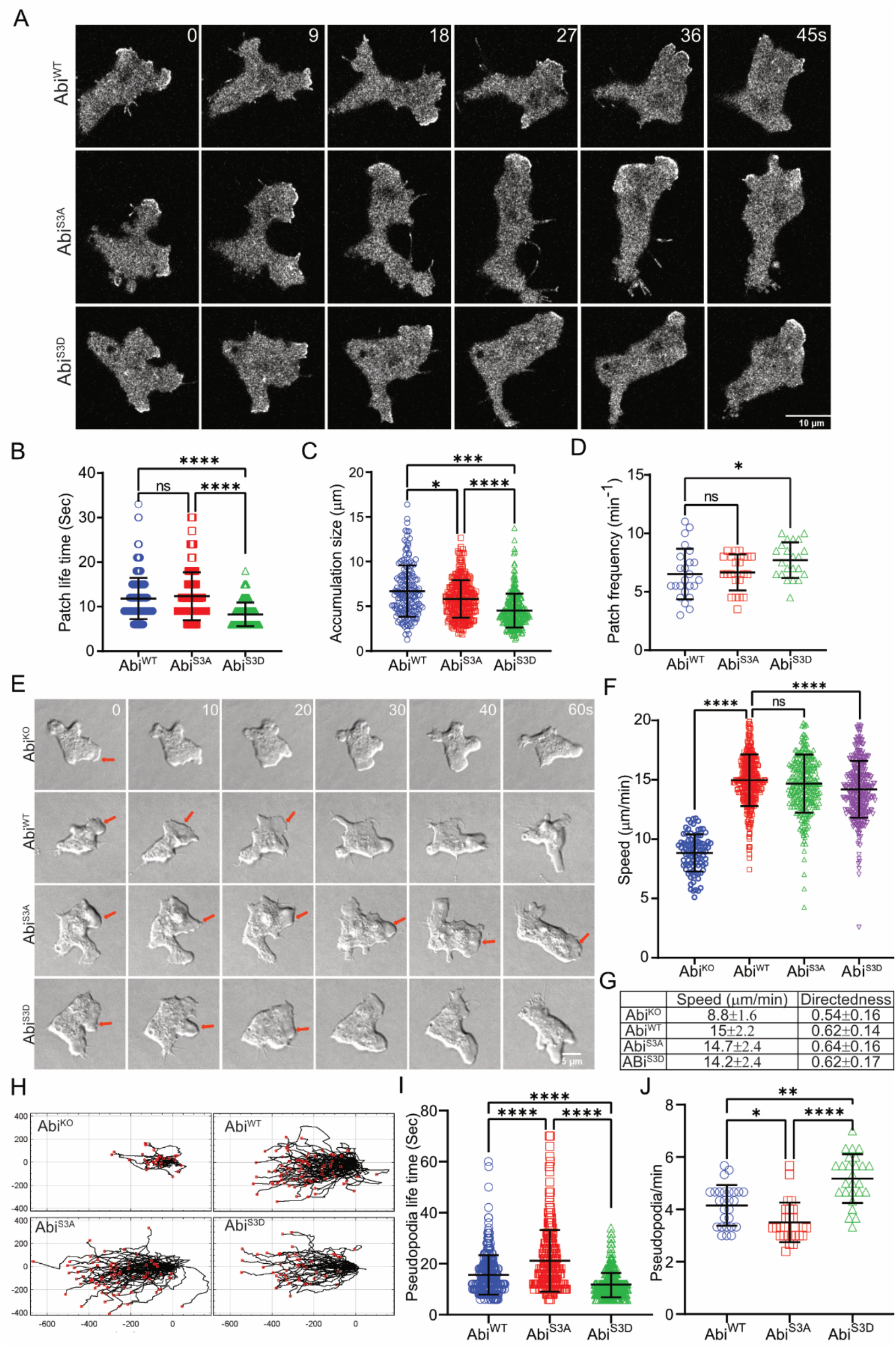
Abi mutants are recruited to pseudopodia and are functional. (A) Subcellular localization of Scar complex. Abi^WT^, Abi^S3A^, and Abi^S3D^ were co-expressed with HSPC300-eGFP in Abi- cells, allowed to migrate under agarose up a folate gradient and examined by AiryScan confocal microscopy at a frame interval of 3 seconds (1f/3s). All show efficient Scar/WAVE complex localization to the pseudopods. (B, C, D) Lifetimes, size, and generation rates of mutant Scar patches. Patch lifetime was measured from the number of frames a patch showed continuous presence of labelled Scar complex, size was measured in ImageJ from individual frames and patch generation frequency was calculated from the number of patches lasting at least 2 frames. Abi^WT^ and Abi^S3A^ patches are less frequent and long-lived compared to Abi^S3D^ (mean ± SD; n > 25 cells over 3 experiments; *p ≤ 0.05, **p ≤0.01, ***p≤ 0.001****p≤ 0.0001, 1-way ANOVA, Dunn’s multiple comparison test). (E) Rescue of pseudopod formation by mutated Abi. Abi- cells were transfected with Abi^WT^, Abi^S3A^, and Abi^S3D^ and allowed to migrate under agarose up a folate gradient while being observed by DIC microscopy at a frame interval of 2 seconds (1f/2s). All rescued cells formed actin pseudopods; those formed by Abi^S3A^ -expressing cells were longer lived and Abi^S3D^ – expressing cells were short lived compared to Abi^WT^. Arrows indicate blebbing in Abi^KO^ and a single pseudopod in rescued cells. Panel (F) shows cell migration speed (mean ± SD; n = 90 Abi-, 327 WT, 296S3A, 347S3D over 3 independent experiments, 1-way ANOVA, Dunn’s multiple comparison test). (G) Trajectories of cells migrating under agarose up a folate gradient. (H) Summary of chemotaxis features of WT and phosphomutants. (I,J) Abi^WT^, Abi^S3A^, and Abi^S3D^ on pseudopod dynamics. Pseudopod lifetime and frequency was measured by counting the number of pseudopods lasting at least 2 frames from DIC videos. Abi^S3A^ yields increased, and Abi^S3D^ demonstrated decreased lifetimes. Abi^S3A^, pseudopods generated less frequently than Abi^WT^ and Abi^S3D^, while Abi^S3D^ generated pseudopods which were more frequent than Abi^WT^ and Abi^S3A^ (mean ± SD; n > 25 cells over 3 experiments; *p ≤ 0.05, **p ≤0.01, ***p≤ 0.001****p≤ 0.0001, 1-way ANOVA, Dunn’s multiple comparison test).

Cells without Abi migrate poorly [20], mainly using blebs rather than actin pseudopods, are very slow (Figure 5E,F&G, Video S8) and less directional (Figure 5G&H). In contrast, Abi^KO^ cells rescued with Abi^WT^, Abi^S3A^ and Abi^S3D^ formed pseudopods that split frequently (Figure 7E, Video S8). The migration speed and directionality of Abi^KO^ was completely rescued by Abi^WT^ or either phospho-mutant Abi with decrease in the speed of cells with Abi^S3D^ (Figure 5 F,G&H). Further, dynamics of pseudopods were altered in mutants (Figure 5 I &J, Video S8). Abi^S3A^ cell pseudopodia lasted longer (21.1±12 seconds, mean±SD) than Abi^WT^ (15.6±7.7 seconds, mean±SD), while Abi^S3D^ pseudopods were shorter-lived (11.8±4.9 seconds, mean±SD). Moreover, pseudopod generation was more frequent in Abi^S3D^ than in Abi^S3A^ (Figure 5J); we interpret this to mean that the smaller, shorter-lived pseudopods in the phosphomimetic mutant were replaced more frequently. These results clearly show that Abi phosphorylation tunes the dynamics of pseudopods after they are formed.

### 3.6 Double Abi/Scar phospho-mutants still rescue migration in Scar-/Abi- cells

Previously, we have shown that phosphorylation of the Scar/WAVE subunit controls pseudopodia dynamics [13, 26]. One hypothesis combining that data, the results in this paper, and the established literature would be that phosphorylation of either Scar/WAVE or Abi is required for the complex to be activated; when one subunit is mutated, phosphorylation on the other is sufficient to compensate.

We therefore tested the combined effects of Scar/WAVE and Abi phosphorylation on the activation of the complex and cell migration. In order to examine the cells’ physiological behaviour accurately and completely, we generated a complex parent. We created a Scar-/Abi- double knockout, and then replaced the endogenous Pir121 with a single copy Pir121-eGFP (Scar-/Abi-/Pir121-eGFP). This yielded a strain in which the Scar/WAVE complex could be followed without tags on either Scar or Abi, or competition with the endogenous proteins, and in which the complex was homogeneously GFP-tagged, giving high signal/noise ratio in microscopy. Co-expression of Scar and Abi, as wild type (Scar^WT^/Abi^WT^), unphosphorylatable (Scar^S8A^/Abi^S3A^) and phosphomimetic (Scar^S8D^/Abi^S3D^) mutants, resulted in consistently normal expression levels of all complex subunits tested, confirming that phosphorylation is not important for complex formation (Figure 6 A&B). We examined the different mutants’ migration in detail, using a folate gradient under agarose. Cells expressing wild-type and mutant Scar/Abi formed pseudopodia with altered dynamics (Figure 6C, Video S9). Scar-/Abi- cells barely form pseudopodia, and those formed are very short-lived, but transfection with all mutants of Scar & Abi rescued pseudopodia formation. Compared with Scar^WT^/Abi^WT^ and Scar^S8D^/Abi^S3D^, Scar^S8A^/Abi^S3A^ formed fewer, more persistent, longer-lived pseudopodia (Figure 6C, Video S9) Scar^WT^/Abi^WT^. Migration speed was also rescued in the Scar-/Abi- cells by expression of wild type or mutant Scar/Abi (Figure 6D&E). There was a marked increase in the speed of Scar^S8A^/Abi^S3A^, compared with Scar^WT^/Abi^WT^ and Scar^S8D^/Abi^S3D^ (Figure 6D&E). Recruitment of the Scar/WAVE complex was rescued in wild type and cells expressing both pairs of phosphomutants (Figure 6F, Video S10), though there were obvious differences. The Scar patches lasted for 11±5.3 sec (mean±SD, n=167) in Scar^WT^/Abi^WT^ (upper panel 1; Figure 6F, Video S10), while Scar^S8A^/Abi^S3A^ patches’ lifetime increased to 16±7.6 sec (mean±SD, n=120) (middle panel; Figure 6F, Video S10 up to frame 36 sec). In contrast, Scar^S8D^/Abi^S3D^ patches were brief (8.7±3.4 sec, mean±SD, n=141) and highly oscillatory (lower panel; Figure 6F, Video S10). Overall, these results suggest that neither Scar nor Abi phosphorylation is required for the Scar/WAVE complex activation. Rather, both sets of phosphorylations are tools that enable the regulation of pseudopodia after they are formed. Each phosphorylation on either Scar/WAVE or Abi shortens the lifetime of the Scar/WAVE complex patches, and therefore the distance covered by each pseudopod.

**Figure 6.**
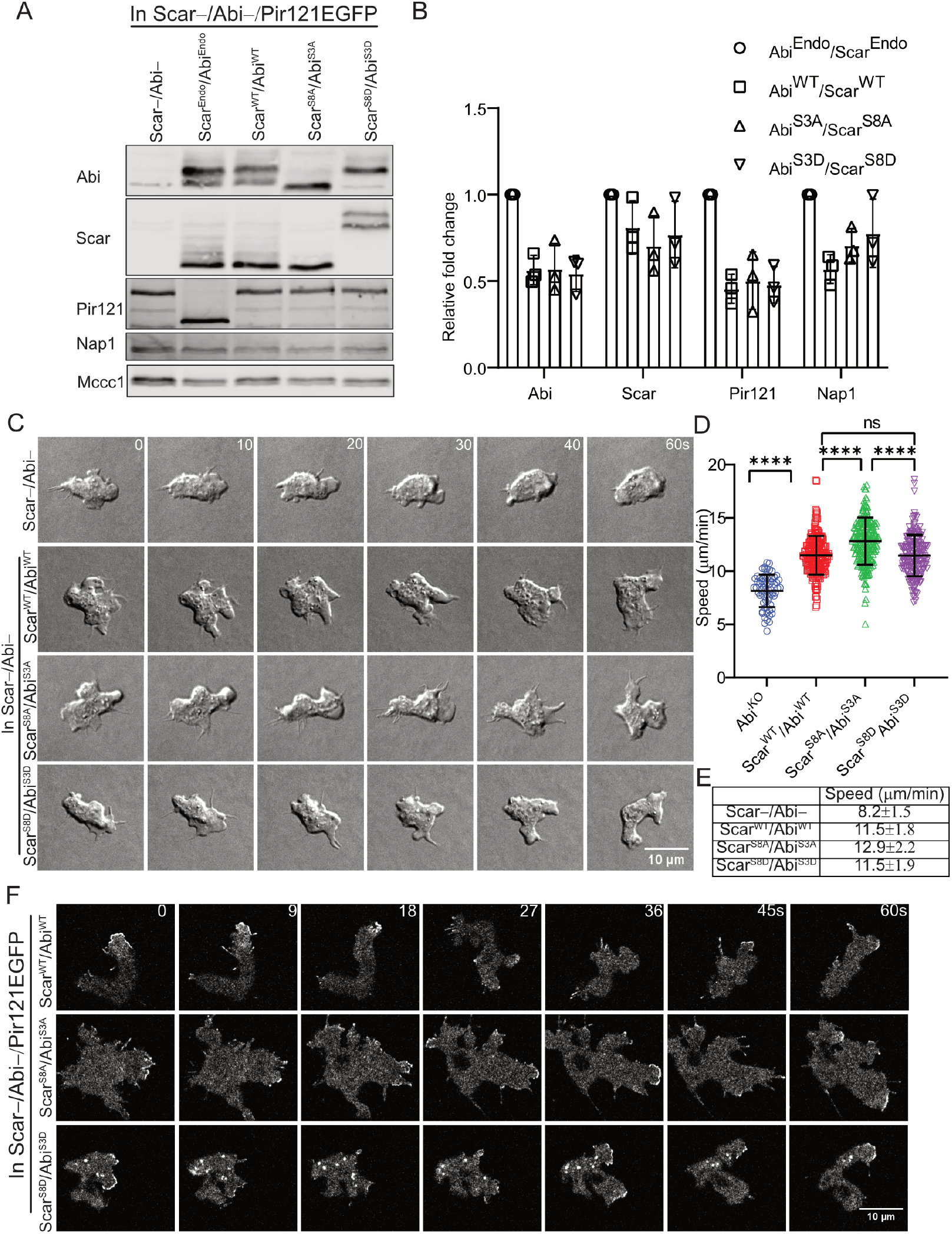
Both Scar and Abi phospho-mutants are recruited to pseudopodia and are functional. (A) Stability of the WT and phosphomutant Scar/WAVE complex. Scar^WT^/Abi^WT^, Scar^S8A^/Abi^S3A^, and Scar^S8D^/Abi^S3D^ were expressed in Scar-/Abi-/Pir121-eGFP cells. Expression of Abi, Scar, Pir121, and Nap1 was compared by western blotting. WT and mutant Scar/Abi expression restored the expression of all subunits. Mccc1 was used as a loading control. (B) Densitometric quantification of protein bands. (C) Rescue of pseudopod formation by WT and phosphomutants of Scar/Abi. Scar-/Abi-/Pir121-eGFP cells were transfected with Scar^WT^/Abi^WT^, Scar^S8A^/Abi^S3A^, and Scar^S8D^/Abi^S3D^ and allowed to migrate under agarose up a folate gradient while being observed by DIC microscopy at a frame interval of 2 seconds (1f/2s). (D) shows speeds (mean ± SD; n = 65 Scar-/Abi-, 193Scar^WT^/Abi^WT^, 233 Scar^S8A^/Abi^S3A^, 195 Scar^S8D^/Abi^S3D^ over 3 independent experiments, 1-way ANOVA, Dunn’s multiple comparison test). (E) Summary of chemotaxis features of WT and phosphomutants. (F) Subcellular localization of the Scar complex. Scar^WT^/Abi^WT^, Scar^S8A^/Abi^S3A^, and Scar^S8D^/Abi^S3D^ were expressed in Scar-/Abi-/Pir121-eGFP cells, which were allowed to migrate under agarose up a folate gradient and were examined by AiryScan confocal microscopy at a frame interval of 3 seconds (1f/3s).

## 4. Discussion

Abi phosphorylation is seen in all species examined. It occurs in the same general region - near the polyproline domain - but the exact sites are not conserved. This is hardly surprising. In both Scar/WAVE and Abi, the polyproline domains are very variable; their existence is conserved between species but their sequence is not. This raises interesting questions about Abi phosphorylation’s mechanisms and biological functions; it is presumably not creating a simple binding site, or it would be conserved in more detail. Our work in this paper shows that it only occurs after the Scar/WAVE complex has been activated, a process that is thought to involve a tightly-folded, autoinhibited complex being converted into an open array with multiple binding sites for other proteins, in particular effectors of actin polymerization like the Arp2/3 complex and VASP. Scar/WAVE itself is phosphorylated in exactly the same way. Thus it is unlikely that phosphorylation of either Scar/WAVE or Abi is fulfilling the role supported by many earlier papers, of causing or allowing the complex to be activated. This was clearly shown by our double mutants - Scar/WAVE complex in which neither Abi nor Scar/WAVE itself can be phosphorylated, is still active. Rather, it is likely that phosphorylation modulates the stability of the open complex, or modulates its affinity for its binding partners.

The precise mechanism by which Abi phosphorylation modulates the Scar/WAVE complex is unclear. This is because so much is yet to be resolved about the complex’s regulation. It is clear that Rac plays a key role, but there are many other interactors that are thought to be important and whose mechanism of action is not known. In particular, the Scar/WAVE complex patches we observe are maintained by positive feedback loops. Phosphorylation could slightly diminish the strength of the positive feedback; a small change in this would effect a large change in the lifetime of the assemblies. An alternative explanation could involve protein stability.

The Scar/WAVE complex is regulated by proteolysis in a complicated way - loss of either of the largest subunits (PIR121 and Nap1) causes a catastrophic loss of Abi, Scar/WAVE itself and HSPC300/Brk1, but loss of Abi does not cause loss of the PIR121/Nap1 dimer. We interpret this to mean there are at least two proteolytic pathways, one that regulates intact complex and PIR121/Nap1, and a far more active one that is specific for the smaller subunits. Phosphorylation of Abi could increase the rate of either pathway and make the activated complex shorter-lived.

Overall, this work does not lessen the importance of Abi phosphorylation - it may not be a key activating step, but we have shown that it changes the extent to which the Scar/WAVE complex promotes actin polymerization, and thus modulates the cell’s movement speed. In an essential process like cell motility, this is a vital role.

## Funding

This work was supported by Cancer Research UK core grant number A17196 and Multidisciplinary Award A20017 to RHI.

## Appendix A

**Figure S1.**
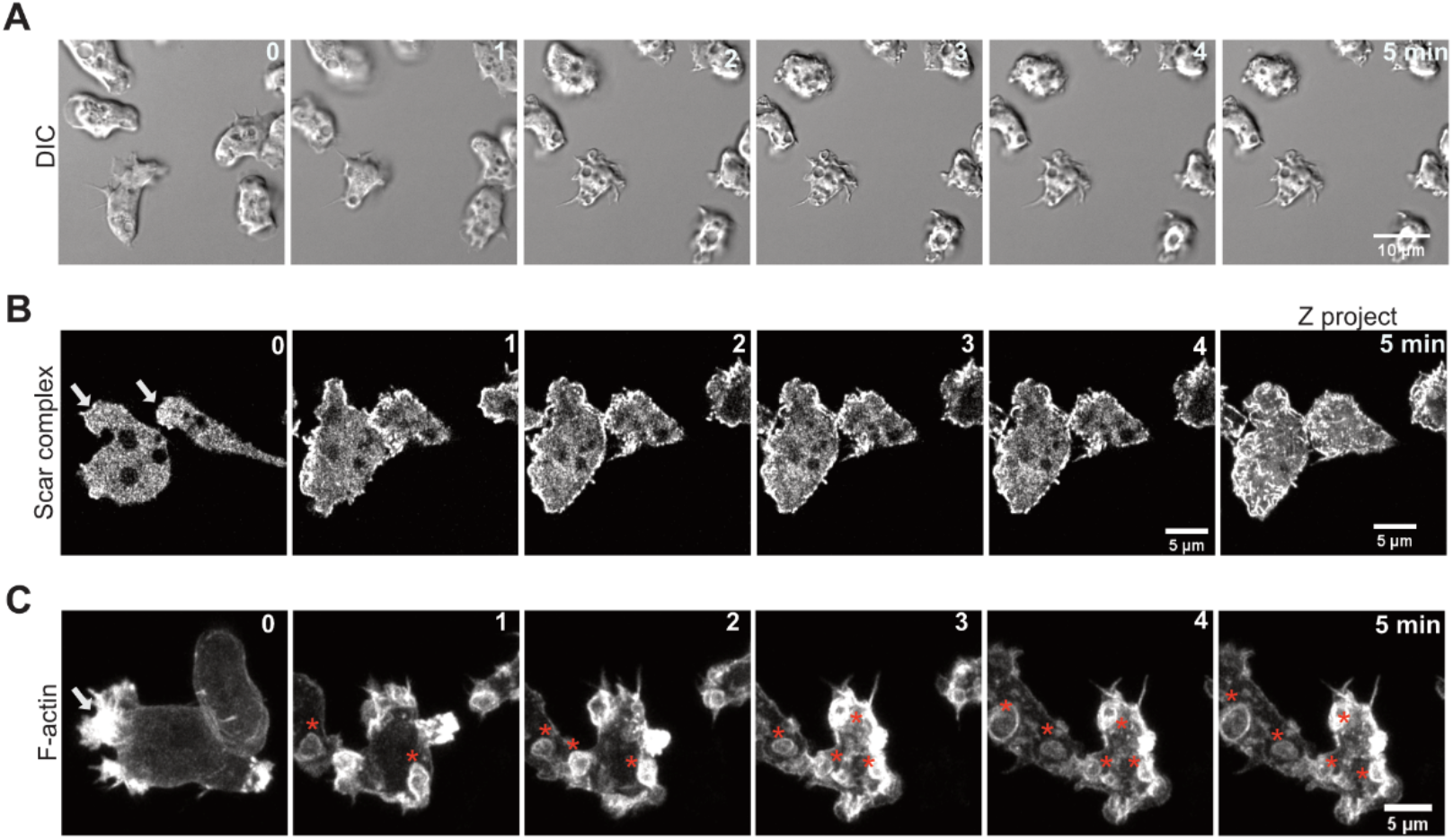
Effect of osmolarity on cell shape, Scar/WAVE and Actin localization. (A) Retraction of pseudopods and shrinkage of cells after addition of sorbitol (0.4M). Cells in glass bottom dishes were imaged by DIC microscopy at a frame interval of 2 seconds (Scale bar =10 mm). (B) Distribution of the Scar/WAVE complex after sorbitol treatment. Cells expressing EGFP-Nap1, marker of the Scar/WAVE complex, were imaged by AiryScan confocal microscopy. 0.4M sorbitol was added during imaging. Sorbitol inhib localization of the Scar/WAVE complex from pseudopodia and causes relocation to the cell cortex (Scale bar= 5 mm). (C) Distribution of actin after sorbitol treatment. Cells expressing lifeact-mRFPmars2 were imaged by AiryScan confocal microscopy. Actin relocates to macropinosomes after addition of sorbitol (Scale bar= 5 mm).

**Figure S2.**
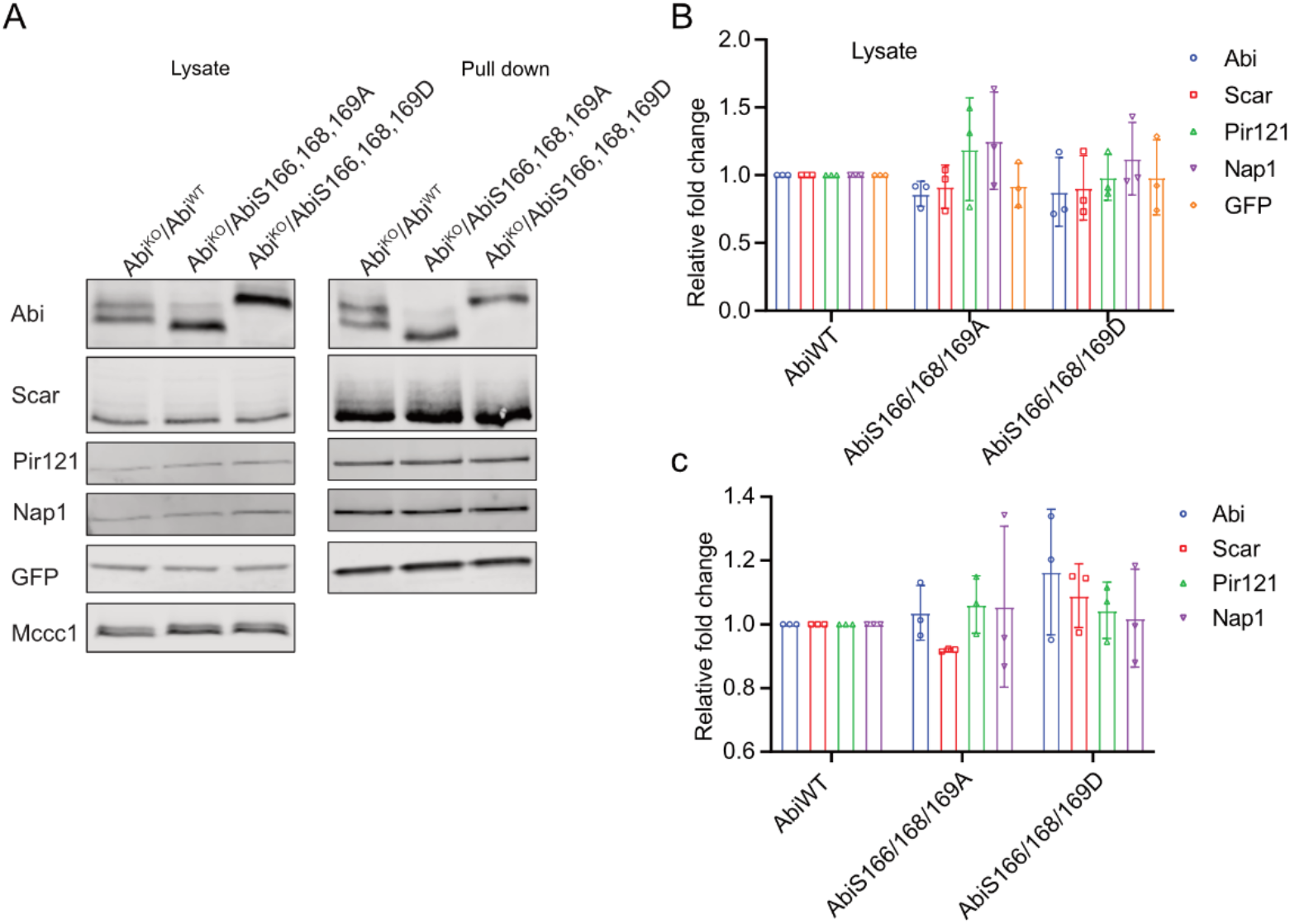
Expression pattern and effect of Abi^WT^, Abi^S3A^, and Abi^S3D^ in the Scar/WAVE complex formation. (A) Abi^WT^, Abi^S3A^, and Abi^S3D^ were co-expressed with HSPC300-eGFP in Abi- cells, and cell lysates were immunoprecipitated using GFP-TRAP. (A) Lysate and pull-down samples were analyzed for the expression of Abi, Scar, Pir121, Nap1 and GFP by western blotting. (B,C) Quantification of western blots shows that Abi^WT^, Abi^S3A^ and Abi^S3D^ formed stable complexes.

**Video S1:** Recruitment of the Scar/WAVE complex after folate treatment. eGFP-Nap1 cells were seeded on Lab-Tek II coverglass chambers and imaged with an AiryScan microscope. Folate (200 μM) was added to the cells undergoing imaging. Filmed at 1 frame/3 seconds, movie shows 10 frames/second. Folate was added after frame 4 (after 12 seconds).

**Video S2:** Recruitment of the Scar/WAVE complex after cAMP treatment. eGFP-Nap1 cells were seeded on Lab-Tek II coverglass chambers and imaged by AiryScan confocal microscopy. cAMP (10 μM) was added to the cells undergoing imaging. Filmed at 1 frame/3 seconds, movie shows 10 frames/second. cAMP was added after frame 4 (after 12 seconds).

**Video S3:** Change in cell morphology after sorbitol treatment. *Dictyostelium* cells were seeded on Lab-Tek II coverglass chambers and imaged by DIC microscopy (60x). Filmed at 1 frame/1 second, movie shows 10 frames/second.

**Video S4:** Recruitment of the Scar/WAVE complex after sorbitol treatment. eGFP-Nap1 cells were seeded on Lab-Tek II coverglass chambers and imaged by AiryScan confocal microscopy. Sorbitol (0.4M) was added to the cells undergoing imaging. Filmed at 1 frame/3 seconds, movie shows 10 frames/second.

**VideoS5:** Recruitment of F-actin after sorbitol treatment. Life-act-eGFP expressing cells were seeded on Lab-Tek II coverglass chambers and imaged by AiryScan confocal microscopy. Sorbitol (0.4M) was added to the cells undergoing imaging. Filmed at 1 frame/3 seconds, movie shows 10 frames/second.

**Video S6:** Recruitment of the Scar/WAVE and Arp2/3 complexes after latrunculin treatment. eGFP-Nap1 cells expressing ArpC2-mRFPmars 2 were seeded on Lab-Tek II coverglass chambers and imaged by AiryScan confocal microscopy. LatrunculinA (5 μM) was added to the cells undergoing imaging. Filmed at 1 frame/3 seconds, movie shows 5 frames/second. Latrunculin was added after frame 1 (after 1 minute).

**Video S7:** Scar complex activation in Abi phosphomutants. Abi^KO^ cells co-expressing HSPC300-eGFP and Abi were allowed to migrate under agarose up folate gradient and Scar complex activation in pseudopods were observed by AiryScan confocal microscopy. Imaged at 1 frame/3 seconds, movie shows 10 frames/second.

**Video S8:** Pseudopod formation in Abi phosphomutants. Abi^KO^ cells expressing Abi^WT^, Abi^S3A^ and Abi^S3D^ were allowed to migrate under agarose up a folate gradient and observed by DIC microscopy. Filmed at 1 frame/2 seconds, movie shows 10 frames/second.

**Video S9:** Scar complex activation in Abi phosphomutants. Scar-/Abi-/Pir121-eGFP cells expressing Scar^WT^/Abi^WT^, Scar^S8A^/Abi^S3A^ and Scar^S8D^/Abi^S3D^ were allowed to migrate under agarose up folate gradient and Scar complex activation in pseudopods were observed by AiryScan confocal microscopy. Imaged at 1 frame/3 seconds, movie shows 10 frames/second.

**Video S10:** Pseudopod formation in Scar and Abi phosphomutants. Scar-/Abi- cells expressing Scar^WT^/Abi^WT^, Scar^S8A^/Abi^S3A^ and Scar^S8D^/Abi^S3D^ were allowed to migrate under agarose up a folate gradient and observed by DIC microscopy. Filmed at 1 frame/2 seconds, movie shows 10 frames/second.

## Appendix B

**Table S1.** Cell lines used in this study.

| No. | Cell line | Given name | Reference |
| --- | --- | --- | --- |
| 1. | Ax3 |  | dictyBase |
| 2. | NC4 |  | dictyBase |
| 3. | Abi- | Ap5b | [20] |
| 4. | eGFP-Nap1 | - | [26] |
| 5. | Scar-/Abi- | SP2 | This study |
| 6. | Scar-/Abi-/Pir121-eGFP | SP21 | This study |
| 7. | Pir121- | SP3 | [4] |
| 8. | Pir121-eGFP WT | SP7 | [4] |
| 9. | Pir121-eGFP A site | SP11 | [4] |

**Table S2.**
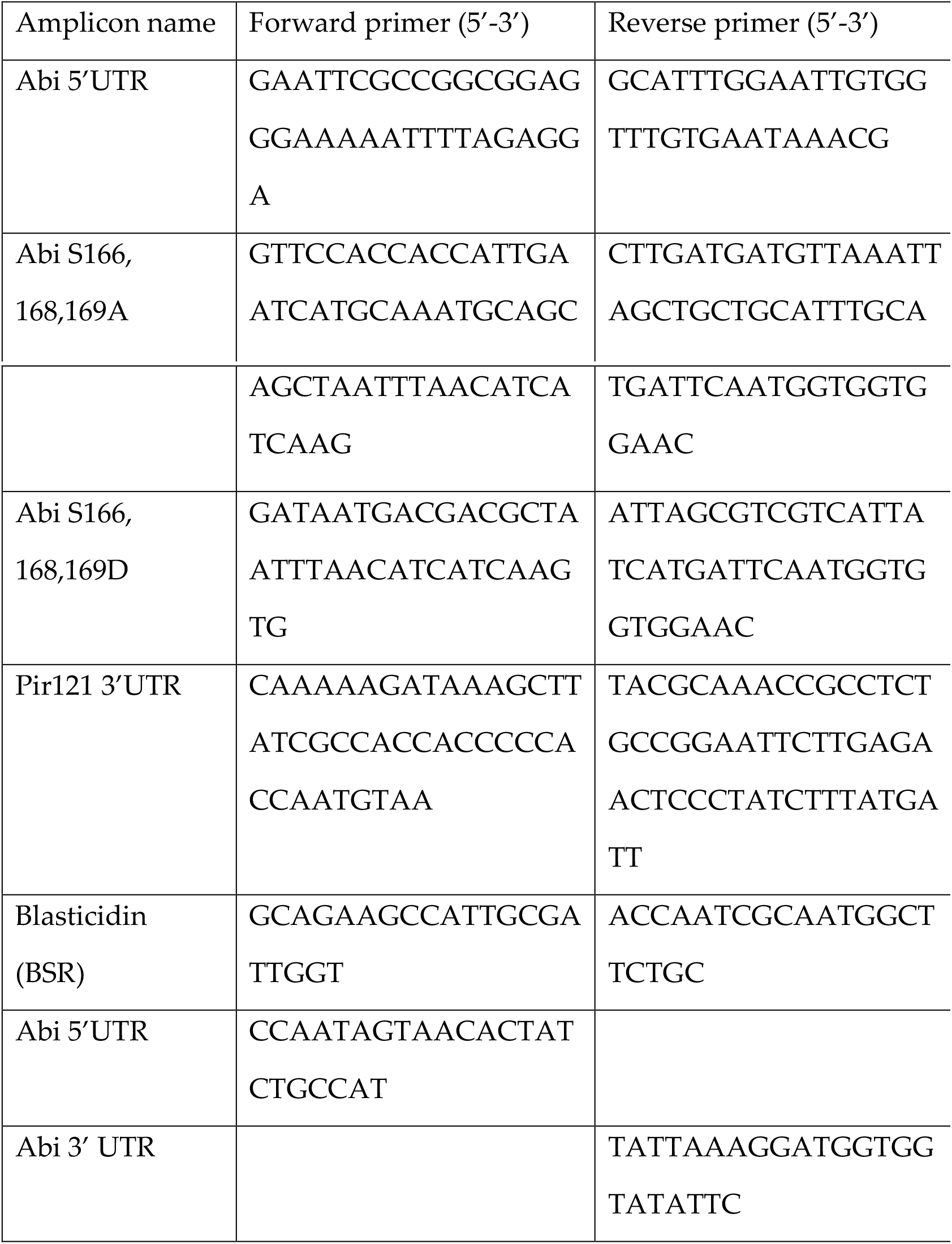
List of primers

**Table S3.** Plasmids created and used in this study

| No. | Name of the plasmid | Gene/s |
| --- | --- | --- |
| 1. | pSP287 | Shuttle vector of Abi WT |
| 2. | pSP318 | Pir121eGFP knock in vector |
| 3. | pSP384 | Abi WT |
| 4. | pSP396 | Abi S166,168,169A (S3A) |
| 5. | pSP427 | Abi S166,168,169D (S3D) |
| 6. | pSP428 | Abi WT and HSPC300-eGFP |
| 7. | pSP429 | Abi S166,168,169A (S3A) and HSPC300-eGFP |
| 8. | pSP430 | Abi S166,168,169D (S3D) and HSPC300-eGFP |
| 9. | pSP487 | ScarWT and AbiWT |
| 10. | pSP488 | ScarS8A and AbiS3A |
| 11. | pSP489 | ScarS8D and AbiS3D |

## Notes

### Competing Interest Statement

The authors have declared no competing interest.

